# The *Nanog* epigenetic remodeling complex was essential to vertebrate mesoderm evolution

**DOI:** 10.1101/2022.09.08.507069

**Authors:** Luke A. Simpson, Darren Crowley, Teri Forey, Helena Acosta, Zoltan Ferjentsik, Jodie Chatfield, Alexander Payne, Benjamin S. Simpson, Catherine Redwood, James E. Dixon, Nadine Holmes, Fei Sang, Ramiro Alberio, Mathew Loose, Andrew D. Johnson

## Abstract

Pluripotency defines the unlimited potential of cells in the primitive ectoderm of vertebrate embryos, from which all adult somatic cells and germ cells are derived. Understanding how the programing of pluripotency evolved has been obscured by the study of early development in models from lower vertebrates in which pluripotency is not conserved. Here we investigated how the axolotl ortholog of the mammalian core pluripotency factor *NANOG*, programs pluripotency during axolotl development to model the tetrapod ancestor from which terrestrial vertebrates evolved. We show that in axolotl primitive ectoderm (animal caps; AC) NANOG synergizes with NODAL activity and the epigenetic modifying enzyme DPY30 to direct the deposition of H3K4me3 in chromatin prior to the waves of transcription required for lineage commitment and developmental progression. We show that the interaction of NANOG and NODAL with DPY30 is required to direct development downstream of pluripotency and this is conserved in axolotls and human. These data demonstrate that the interaction of NANOG and NODAL signaling represents the basal state of vertebrate pluripotency.

## Main

Pluripotency defines the competency of early embryonic cells to differentiate into any somatic cell type or primordial germ cells (PGCs). Therefore, pluripotency underpins the embryo’s ability to form a functional animal. How the programing of pluripotency has evolved is unclear in part because many non-mammalian vertebrate models do not display pluripotency during early development^1-3^. Axolotls model the development of the tetrapod ancestor; indeed, the AC of axolotls can form all three primary germ layers and germ cells in response to inductive signals^4^. Given the role of the transcription factor NANOG in programming pluripotency in pre-implantation mammalian embryos^5^, here we investigated how NANOG programs pluripotency in axolotl development. We show that in axolotl ACs, NANOG synergizes with NODAL activity and the epigenetic modifying enzyme DPY30 to direct the deposition of H3K4me3 to facilitate lineage commitment and developmental progression. We show that the interaction of NANOG, NODAL and DPY30 is required to direct development downstream of pluripotency and this mechanism is likely conserved between axolotls and human. These data demonstrate that the interaction of NANOG and NODAL signalling are key to establishing pluripotency in vertebrates.

### Nanog is required for the development of axolotl embryos

RNA sequencing (RNA-seq) on axolotl AC explanted at mid-gastrula stage (stage 10.5) showed that the pluripotency genes *Nanog* and *Pou5f1 (or Oct4)* were highly expressed (Fig.1 a-b). Integration of this RNA sequencing and sc-RNASeq datasets^6-8^ from stage matched *Xenopus* tropicalis AC, peri-gastrulation pig and human epiblast cells, showed similar expression profiles of pluripotency factors *SOX2 and LIN28A* in all four organisms. However, only axolotl, pig and human pluripotent cells showed similar expression profiles for the pluripotency genes *NANOG, OCT4 and PRDM14* (Fig.1 b). Other pluripotency genes, such as *TFCP2L1* and *OTX2*, showed similar expression between human-axolotl and pig-axolotl, respectively. Interestingly, few genes were expressed in mammalian and *Xenopus* cell populations but not axolotl. Given that *NANOG* and *OCT4* are not conserved in the *Xenopus* genome^9^, differences observed here may reflect divergence of the pluripotency gene regulatory network (pGRN) in frogs from a conserved vertebrate state.

**Figure 1.**
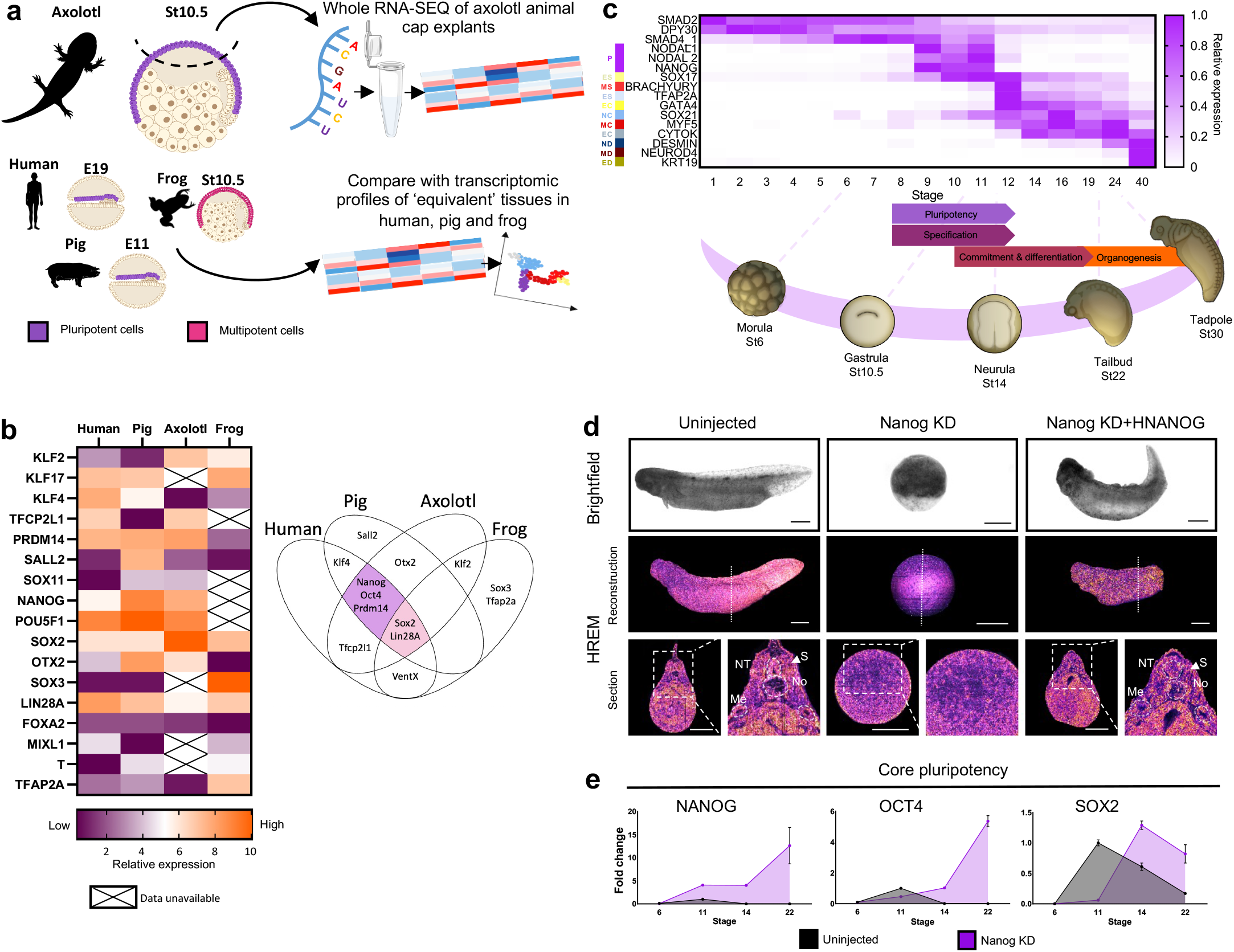
AC pluripotency resembles that of mammals and is under the control of Nanog. **a**, Outline of transcriptomic data collection and cross-species comparison (n=15×3). **b**, Key pluripotency gene expression in the peri-gastrula primitive ectoderm of axolotl, frogs and mammals **c**, Overview of axolotl early development. Pluripotency (P), endoderm specification (general) (ES), mesoderm specification (general) (MS), ectoderm specification (general) (EcS), mesoderm commitment (MC), endoderm commitment (EC), ectoderm commitment (EcC), mesoderm differentiation (MD), endoderm differentiation (ED), Neural differentiation (ND). **d**, Brightfield images and HREM reconstructions of uninjected and Nanog depleted and hNanog rescued embryos (n=25 and 2 respectively). Dotted line marks plane of section reconstruction. Dashed lines highlight: somites (S), neural tube (NT) notochord (No), mesonephric ducts (Me). Data fromk Jiang *et al*^15^. Scale bar, 1mm. **e**, Expression of core pluripotency genes at different developmental stages with and without Nanog KD (n=10×3).

Mouse pluripotent cells are classified as either naïve or primed, defined by their developmental potential and their dependence on nodal signalling^10^. In axolotl, both *Nodal* and *Nanog* are activated between stages 8 and 9 in the AC following zygotic genome activation (ZGA)^11,12^ (Fig. 1c). Moreover, there is no stage at which *Nanog* is expressed in the absence of *Nodal*, suggesting that their co-expression represents basal pluripotency (Fig. 1c).

Antisense morpholinos (MO) to knockdown (KD) NANOG expression, either by inhibiting translation or splicing (Fig. 1d; Extended Data Fig. 1a-f), result in the complete arrest of development at stage 9, prior to the onset of gastrulation. Normal development was rescued by injection of 100pg human *Nanog* (*hNanog*) RNA (Fig. 1d; Extended Data Fig. 1d&f) demonstrating specificity. High-resolution episcopic microscopy (HREM) of NANOG morphants showed no evidence of either involution or ingression, which characterize axolotl gastrulation^13^ (Fig. 1d, Extended Data Fig. 1e). Morphants maintained this arrested morphology for weeks, surviving beyond Tailbud stages in matched controls. It has been suggested that *Xenopus Ventx1/2* is functionally equivalent to NANOG, but *Ventx1/2* KD does not arrest development at blastula stage^14^, suggesting that the two genes are not functionally equivalent. Gene expression analysis of NANOG morphants demonstrated the persistent expression of *Nanog, Oct4* and *Sox2* (Fig. 1e), indicating that morphants failed to extinguish pluripotent gene expression long after uninjected siblings completed gastrulation. This suggests that NANOG is critical to extinction of pluripotency during embryogenesis.

### Germ-layer commitment requires NANOG activity

Genome-wide effects of NANOG KD on cell fate decisions were measured during gastrulation (stage 10.5) or tailbud (stage 22) stages (Fig. 2a-b)^15^. Gastrula cell-specific markers were either up or downregulated and many zygotic genes were activated at normal or elevated levels showing NANOG is not required for ZGA^16^. By tailbud, however, pluripotency and mesendoderm specification genes showed higher expression in morphants (Fig. 2a and Extended Data Fig. 1g). These genes are normally downregulated by this stage^15^, suggesting NANOG is required to progress beyond lineage specification. Accordingly, genes involved in mesoderm or ectoderm commitment, including *Hox* genes, were downregulated, while endodermal commitment genes increased (Fig. 2a-b and Extended Data Fig. 1g). Gene-set enrichment analysis (GSEA) (Extended Data Fig. 2a) showed enrichment of lateral plate, notochord, cardiac mesoderm, neural tube, neural crest, and epidermal progenitor markers among downregulated genes in NANOG morphants. Moreover, stage 28 morphants showed loss of differentiation markers (Extended Data Fig. 1h), suggesting NANOG depletion causes embryos to form endoderm-like tissue that does not differentiate. These findings suggest NANOG is required to coordinate lineage specification and commitment. Depletion, therefore, ablates the successive waves of gene expression that drive embryogenesis. Further, results indicate that embryos default to an endoderm-like state (Fig. 2c).

**Figure 2.**
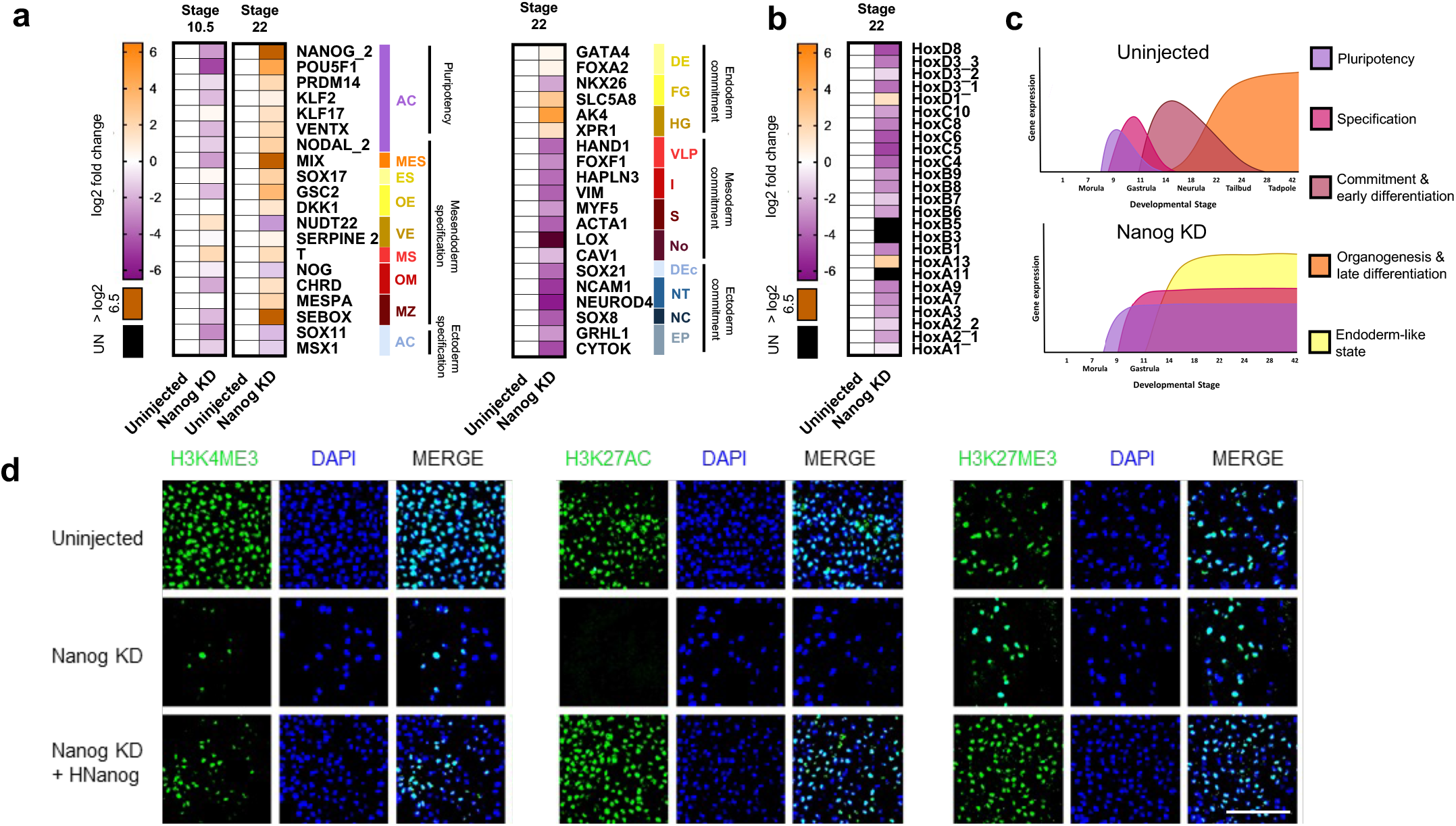
Nanog is required for waves of embryogenesis. **a**, Differential gene expression of cell-type marker genes at stages 10.5 and 22 in uninjected (n=7 and 6, respectively) and Nanog depleted (n=6 and 3, respectively) embryos. Black indicates no detectable expression. Animal cap (AC), mesendoderm specification (general) (MES), endoderm specification (general) (ES), organiser endoderm (OE), vegetal endoderm (VE), mesoderm specification (general) (MS), organiser mesoderm (OM), marginal zone (MZ), definitive endoderm (general) (DE), foregut (FG), hindgut (HG), ventral-lateral plate (VLP), intermediate mesoderm (I), somite (S), notochord (No), definitive ectoderm (general) (Dec), neural tube (NT), neural crest (NC), epidermal progenitors (EP). **b**, Expression of Hox genes in response to Nanog KD at stage 22. **c**, Diagrammatic representation of the effect of Nanog depletion which prevents the sequential waves of gene expression. **d**, Uninjected, Nanog depleted and hNanog rescued AC explants cultured to equivalent stage 14 and stained for H3K4me3, H3K27ac, H3K27me3 and DAPI (n=3). Scale bar, 60µm.

The developmental potential of NANOG depleted AC was then addressed directly using explants (Extended Data Fig. 3a). Caps normally differentiate into epidermis, marked by *Grhl1* and *Cytok*, and down-regulation of pluripotency markers. Morphant caps, however, failed to produce epidermis and maintained high *Nanog* and *Oct4* levels (Extended Data, Fig. 3b). In AC, 1pg of RNA encoding activin induces mesoderm, marked by elongation and *brachyury* expression^11^. In contrast, 200fg of RNA is insufficient to induce mesoderm (Extended Data Fig. 3d). Remarkably, when combined with NANOG depletion, caps expressing sub-mesodermal (200fg) doses of activin expressed markers for endoderm, though not mesoderm. 1pg of activin induced high levels of endoderm markers, also without induction of brachyury (Extended Data Fig. 3d), a response normally attributed to very high levels of activin. These data suggest that AC depleted of NANOG are hyper-sensitized to TGF-β signalling. We posit, therefore, that NANOG may act as a rheostat of SMAD2 activity, as has been shown for SMAD1/5 in mouse ESC^13^, which in AC modulates the response to NODAL signalling levels (Extended Data, Fig. 3f). This aligns with the inability of *Nanog* null mouse embryos to complete epiblast specification^5,17^.

**Figure 3.**
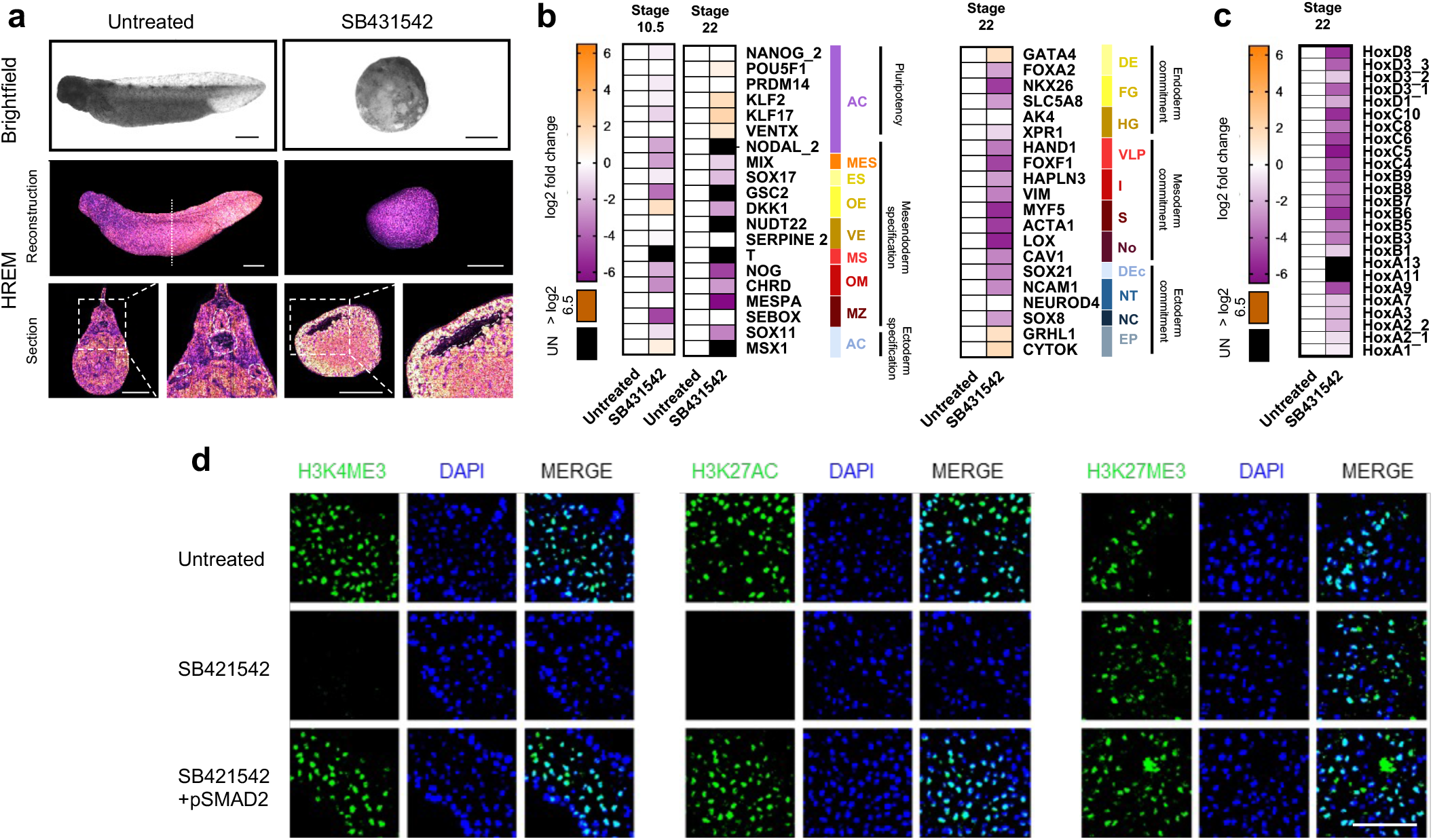
Nodal signalling programs early germ layer specification. **a**, Brightfield and HREM images of uninjected and SB431542 treated embryos. Dotted line marks plane of section reconstruction. Somites (S), neural tube (NT) Notochord (No), Mesonephric ducts (Me), Blastocoel (B) (n=25 and 2 respectively). Scale bar, 1mm. **b**, Gene expression of cell-type marker genes at stages 10.5 and 22 in uninjected (n=7 and 6, respectively) and SB431542 (n=6 and 6, respectively) treated embryos. Black indicates no detectable expression. Animal cap (AC), mesendoderm specification (general) (MES), endoderm specification (general) (ES), organiser endoderm (OE), vegetal endoderm (VE), mesoderm specification (general) (MS), organiser mesoderm (OM), marginal zone (MZ), definitive endoderm (general) (DE), foregut (FG), hindgut (HG), ventral-lateral plate (VLP), intermediate mesoderm (I), somite (S), notochord (No), definitive ectoderm (general) (Dec), neural tube (NT), neural crest (NC), epidermal progenitors (EP) **c**, Gene expression of Hox genes in SB431542 treated embryos at stage 22. **d**, Uninjected, SB431542 treated and SMAD2 rescued AC explants cultured to equivalent stage 14 and stained for H3K4me3, H3K27ac, H3K27me3 and DAPI (n=3). Scale bar, 60µm.

In amphibians, foregut derives from organiser endoderm while hindgut is produced from the vegetal hemisphere^18^. We found that vegetal hemisphere explants at late stages express *Endodermin*, a marker of definitive endoderm (Extended Data Fig. 4h). However, we have not detected *Endodermin* in morphant ACs expressing early endoderm markers, even after supplementation with exogenous activin, though these caps will express *C8b*, an endoderm commitment marker expressed after gastrulation (Extended Data Fig. 3e). This suggests that morphant primitive ectoderm arrests endoderm development at the progenitor stage. Transcriptomes show expression of foregut and hindgut markers in control embryos, but only hindgut markers in morphants (Extended Data Fig. 3e). We surmise, therefore, that hindgut endoderm in morphant embryos is derived from the vegetal pole, which does not express *Nanog*^12^, while anterior endoderm development from animal hemispheres is arrested by *Nanog* depletion.

### NANOG and NODAL activity are required for gastrulation in axolotl

In human ESC, *nanog* synergises with *nodal* signals to deposit H3K4me3 on promoters of transcriptionally active genes^19^. We investigated if this is conserved in axolotl AC. We first investigated when axolotl genomes acquire epigenetic organisation. Western blotting determined that H3K4me3, H3K27me3 and H3K27ac are gradually acquired between stages 8.5 and 10, around the same time *Nanog* is first expressed (Extended data Fig. 4a). Extracts from animal or vegetal poles at stage 10.5 showed, surprisingly, that epigenetic marks were only detectable in AC (Extended data Fig. 4b). ACs from embryos at stage 10.5 were then used for immuno-staining with antibodies to each mark, showing they are readily detectable in nuclei (Fig. 2d). However, expression of H3K4me3 and H3K27Ac, in NANOG morphant caps at the same stage, was no longer detectable with H3K27me3 unaffected. The marks were rescued using *hNanog* RNA (100pg). We confirmed absence of H3K4me3 and H3K27Ac in morphant caps using Western blotting (Extended data Fig. 4c). We also showed that native H3, H3K36me3 and phospho-POLII, which demarcates coding regions of actively transcribing genes, were also unaffected by NANOG depletion (Extended data Fig. 4d&f). This affirms that NANOG morphant ACs are transcriptionally active, but morphant transcription is non-productive and cannot drive embryogenesis. Our data align with human ESC in respect of the role of *nanog* in targeting active transcription marks in pluripotent chromatin^19^, suggesting conservation in vertebrate evolution.

*Nanog*’s epigenetic regulation in hESC depends on interaction with *smad2/3*, downstream of active *nodal* signalling^19^, so we asked if this functions in pluripotent AC. We previously demonstrated that axolotl mesoderm induction is mediated by *activin/nodal* signalling and axolotls share a conserved mesoderm GRN with mammals^11^. Moreover, the phenotype from nodal signalling inhibition using SB431542 (SB) resembles the axolotl NANOG morphant phenotype shown above, which arrests development prior to gastrulation. HREM confirmed the absence of morphological development following SB exposure (Fig 3a). Transcriptomes of SB treated embryos showed down-regulation of mesendodermal specification genes (Fig. 3b-c; Extended data Figs. 2b&5a-b). However, NODAL inhibition, while completely blocking mesendoderm specification, had little effect on pluripotency genes, in contrast to NANOG KD. Like *Nanog* morphants, however, stage 22 SB treated embryos lacked mesodermal commitment markers, as well as neurectoderm markers, confirmed by GSEA (Extended data 6b). SB up-regulated genes were enriched for markers of epidermal cell-types, suggesting that ectoderm defaults to epidermal development absent nodal signalling. We did not observe an effect on definitive endoderm, nor hindgut markers; however, markers of foregut were downregulated. This suggests that hindgut progenitor commitment is independent of activin/nodal signalling. We asked if expression of mature endodermal markers in SB treated embryos was due to NODAL independent expression from the vegetal pole. We treated vegetal pole explants with SB and analysed endodermal gene expression (Extended data Fig. 5d). Indeed, endodermal marker expression from vegetal poles was unaffected by SB, indicating that hindgut endoderm develops independent of NODAL signalling. Interestingly, endoderm is the most ancient metazoan germ layer^20,21^, and may have predated the evolution of *nodal* signalling^22^, suggesting that vegetal pole endoderm reflects an ancient metazoan state.

### NANOG and NODAL synergize with DPY30 to remodel the epigenome post-ZGA

We next asked if the effects of SB treatment could be due to epigenetic effects^19^. Immunostaining confirmed that nodal signalling inhibition results in loss of H3K4me3 and H3K27ac but does not reduce H3K27me3 (Fig. 3d), similar to NANOG depletion. Importantly, when we expressed a sub-mesodermal inducing level (1pg) of RNA encoding a constitutively active (signalling independent) SMAD2 variant^23^ in SB treated caps we rescued these marks (Fig. 3d), demonstrating their dependence on SMAD2/3, which we validated with Western blotting (Extended Data Fig. 4c). These data strongly suggest that the complex of NANOG and SMAD2/3 that regulates transcription activating epigenetic mark deposition in hESC^19^ is conserved in axolotl pluripotent cells.

Contrasting with mammalian *NANOG*s, axolotl *Nanog* encodes a monomer^12^. To test if monomeric NANOG interacts with SMAD2 we used luciferase complementation and showed axolotl NANOG interacts with SMAD2 in mammalian cells (Extended data, Fig. 5e). We then knocked down the epigenetic modifying enzyme DPY30 (Extended data, Fig. 7a-i), which catalyses tri-methylation of H3K4 residues as part of the COMPASS complex and interacts with *nanog* and *smad2/3* in hESC^19^. Unlike the NANOG KD or SB phenotypes, however, DPY30 KD embryos arrested post-gastrulation (Extended Data Fig. 7a&b). HREM showed these morphants lacked a notochord, somites or other mesodermal structures, had a multi-layered AC and rudiments of a neural tube. Remarkably, mouse embryos depleted of DPY30 also show neural development in the absence of mesoderm^19^, suggesting morphological consequences of DPY30 KD are similar in axolotl and mammals. Embryos resembling wild type were rescued with 95% efficiency by 200pg RNA encoding human *DPY30*, demonstrating specificity. Moreover, H3K4me3 became undetectable in ACs after DPY30 depletion, and H3K27ac was diminished shown by immunostaining and Western blotting (Extended Data Fig. 4c&6J). Given the H3K4 tri-methyltransferase activity of DPY30, this suggests that loss of H3K27ac may be an indirect effect of NANOG/DPY30/SMAD2/3 (NDS) depletion. Transcriptomes confirmed DPY30 morphants show reduced expression of pluripotency and lineage specification genes in gastrulae but failed to transcriptionally silence these genes (Extended Data Fig. 7c), also resembling NANOG morphants. DPY30 mutants showed severe reduction in expression of mesodermal and foregut endoderm markers and reduced Hox gene expression (Extended Data Fig. 7d). Hindgut endoderm and neural plate markers were not, however, downregulated. GSEA confirmed that genes downregulated in response to DPY30 KD were significantly enriched for mesodermal tissue markers (Extended Data Fig. 8). By contrast, the most up-regulated genes were enriched for blastula and organiser endoderm markers. Additionally, markers of late mesodermal differentiation were not observed in the morphants (Extended Data, Fig. 7e&f), while endoderm and ectoderm markers were much less affected. Together, these results suggest that DPY30 may regulate the formation of the same tissues in mammals and urodeles and the overlapping effects of NANOG, SB and DPY30 suggest that function of the NDS complex has been conserved through vertebrate evolution.

To test for overlap between effects of NODAL signalling inhibition, or expression of NANOG or DPY30, we compared transcriptomes from embryos under each treatment regime (Extended Data Fig.9, Fig.4a). We identified 1508 ‘early activated’ genes whose expression is usually reduced between stage 10.5 and 22 that are significantly up regulated in NANOG and DPY30 morphants, including pluripotency and early specification genes. We also observed overlap in genes normally up-regulated between stage 10.5 and 22 that are instead down-regulated by each treatment. GSEA showed down-regulated genes were enriched for markers of mesoderm and neural tissues (Fig. 4b). Remarkably, however, while mesendodermal specification genes are downregulated by NODAL signalling inhibition they are upregulated by NANOG or DPY30 KD at later stages, suggesting SMAD2/3 can act independent of interaction with NANOG in early specification events. These transcriptional effects were then correlated with changes in H3K4me3 levels using ChIP-qPCR (Fig. 4c&d, Extended Data Fig. 10). ChIP revealed H3K4me3 levels on promoters of *eef1a1* or *cytok* were unaffected in stage 10.5 caps under each experimental condition (Extended Data Fig. 10a), suggesting expression is independent of the NDS complex. However, H3K4me3 was reduced under all three conditions on the *Nanog* and *Nodal1* promoters (Fig. 4c). Given that these genes first express at ZGA, we posit that their activation is mediated by maternal factors not dependent on NDS, but that the H3K4me3 mark may be required to integrate their expression into the pGRN that extinguishes their expression after gastrulation to initiate the subsequent waves of gene expression that drive development.

**Figure 4.**
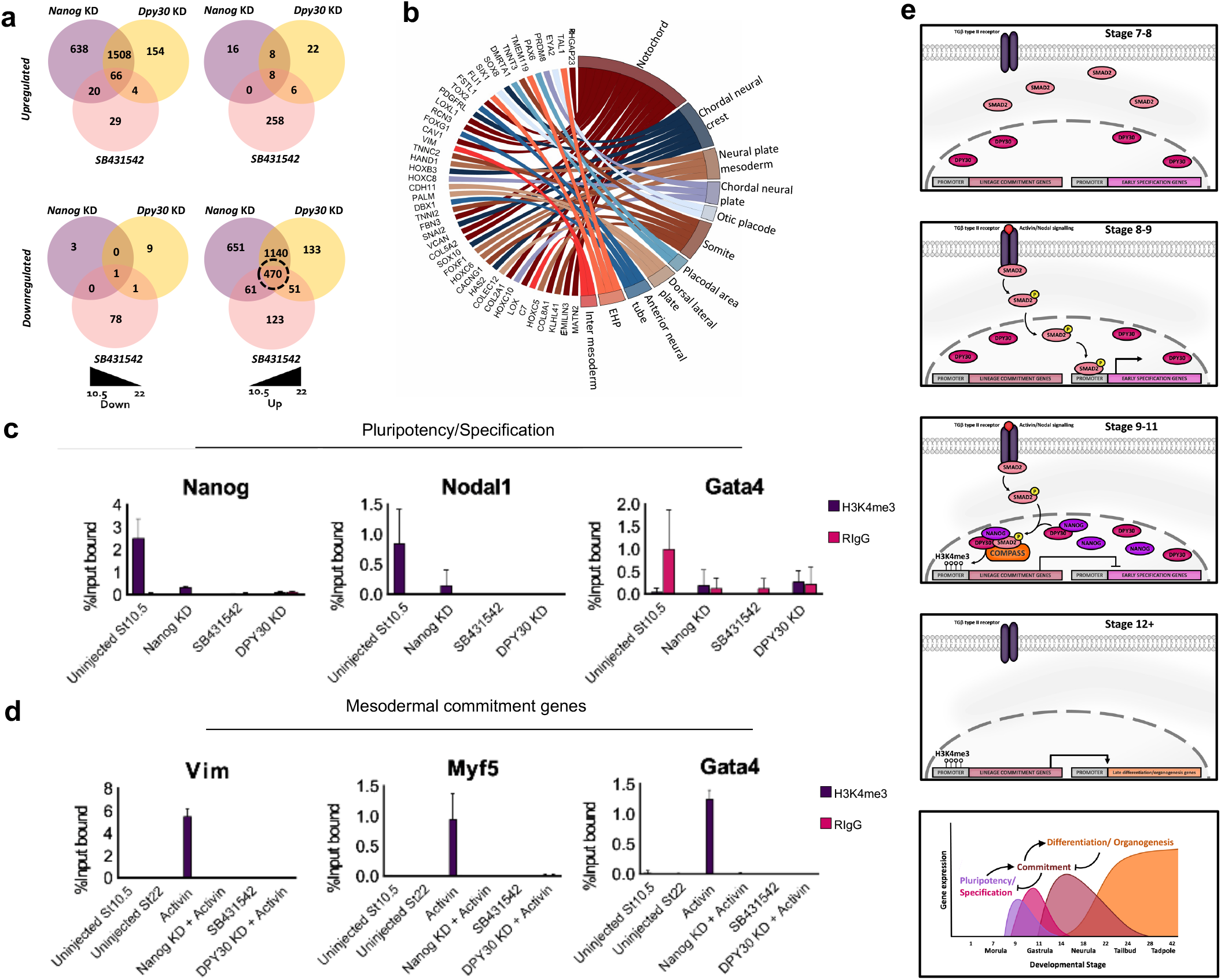
Nanog gene target loci lack H3K4me3. **a**, Venn diagram of overlapping differentially expressed genes in Nanog, DPY30 and Nodal depleted embryos at stages 10.5 and 22. **b**, Enrichment of amphibian cell-type specific markers in overlapping downregulated genes. **c**, H3K4me3 ChIP of stage 10.5 uninjected, Nanog KD, DPY30 KD and SB431542 treated caps followed by qPCR using probes directed at gene promoter regions (n=50 per experimental condition). **d**, H3K4me3 ChIP-qPCR of equivalent stage 10.5 and 22 Uninjected caps, stage 22 Activin treated caps with and without Nanog and DPY30 depletion as well as SB431542 treated caps (n=50 per experimental condition). **e**, Schematics of hypothesis outline.

We then tested effects of each experimental regime on mesoderm induction by performing ChIP on stage 22 caps that had also been co-injected with 1pg of activin RNA along with either the NANOG or *DPY30* MO. H3K4me3 levels were then compared with caps at the same stage that were untreated or exposed to SB. In addition, we included untreated caps at stage 10.5 to determine if the H3K4me3 mark is induced by activin or laid down prior to mesoderm inducing signals (Fig. 4d, Extended data Fig.10b). ChIP revealed that activin induced deposition of H3K4me3 on promoters of mesodermal commitment genes *Myf5 and Vimentin*, and this was prevented by either NANOG or DPY30 KD. This mark was also absent in uninjected and SB treated ACs correlating with the absence of expression. Both *Nodal1* and *Gata4* promoters also show reduced enrichment of H3K4me3 in response to depletion of *Nanog* and *DPY30* (Extended Data Fig. 10b, Fig. 4d). Like uninjected caps, those treated with SB also showed no enrichment of H3K4me3. Whilst expression of these genes is not present in uninjected or SB treated caps, they were activated after either NANOG or DPY30 depletion, and their expression was not extinguished post-gastrulation.

Combined, these data reinforce the concept that NANOG and SMAD2/3 regulation of H3K4me3 is required for mesodermal commitment. Furthermore, this suggests SMAD2/3 regulates transcription of some early genes independently of NANOG. However, both NODAL and NANOG activities are required to regulate the sequential waves of embryonic gene expression in vertebrate development. We posit the following model (Fig. 4e) describing the events from fertilisation to the acquisition of pluripotency. SMAD2 and DPY30 are maternally inherited molecules, highly expressed prior to ZGA in the stage 8 blastula. Around stages 8-11, zygotic *Nodal* and *Nanog* expression commences, the former facilitating phosphorylation of SMAD2, enabling translocation into the nucleus and activation of *Nodal* target genes. Phosphorylated SMAD2 is also able to form a complex *de novo* with NANOG and DPY30/COMPASS which is required to prime mesodermal commitment gene promoters either directly, or via genes acting upstream, through the deposition of H3K4me3. The activation of lineage commitment genes from stage 11 onwards is absolutely required to extinguish expression of pluripotency and early specification specific genes.

Based on our findings we postulate that *NANOG* evolution was integral to the emergence of complex vertebrate mesoderm, as well as the foregut endoderm, defining basic distinctions between vertebrates and more primitive species. In our view, *NANOG* was absolutely required to elaborate the potential of pluripotent cells into the complex tissue types characteristic of vertebrates, making Nanog the defining feature enabling vertebrate evolution.

## Supplementary Information

is available for this paper.

## Acknowledgements

This project was funded by grant from UK MRC [MR/N020979/1] (to AJ).

## Author Contributions

A.D.J conceived of the project with input from M.L and R.A. L.S, D.C, H.A, Z.F, J.C, J.D and N.H performed experiments. T.F, M.L, F.S, B.S, A.P and L.S performed computational analysis. L.S prepared figures. The manuscript was written by L.S in conjunction with A.D.J, M.L and R.A. The manuscript was finalised after review by all authors.

## Methods

### Axolotl embryos and explants

Procedures involving animals have been approved by the Animal Welfare Ethics Committee Review Board, University of Nottingham. Embryos were collected following matings as described previously^24^. For microinjection, embryos were manually de-jellied and cultured in 1× modified Barth’s solution (MBS) with 4% Ficoll (Sigma). Embryos were staged as previously^4,11^. From stage 7 onwards, embryos were maintained in 0.2× MBS, and dissected explants were maintained in 0.7×Marc’s modified ringers solution (MMR). Culture solutions were supplemented with antibiotics (50 µg/ml penicillin and streptomycin, and 50 µg/ml kanamycin), 100 µg/ml Ampicillin and 50 µg/ml fungizone).

### Morpholino and RNA microinjections

Morpholino oligonucleotides (GeneTools, LLC,OR) were designed to block translation of target proteins. Intron/exon boundaries were predicted by homology. Sequences were obtained from our axolotl genomic resource (website URL not finalised)^25^, or via the axolotl genome website^26^. The morpholino sequences used were as follows: Translation MO: *Nanog*, 5′- GGTCAATCCAAAAGCTCCTCCTAAG- 3′; Splice MO: *Nanog* 5′-GGCAGGACTGAAACAAAACGAAGAC-3′; Translation MO: DPY30 5′-ATGCTTTGTTCCGACTCCATTGTGA -3′ and 5′- TTATCGTAGCCCGTCACTCCAGCTC-3′. A nonspecific morpholino was injected in each experiment at equivalent levels to the specific splice morpholino combinations: MO: Control, 5′-GGATTTCAAGGTTGTTTACCTGCCG-3′. Each morpholino experiment was repeated at least three times, and the efficacy of the splice moropholinos was tested by PCR in each experiment using primers detailed in table 1. The Activin-nodal inhibitor SB431542 (Sigma) was solubilised in dimethyl sulfoxide and used at a final concentration of 150 μM. In vitro transcription and microinjection mRNAs for microinjection were synthesised using mMessage mMachine (Ambion) from plasmids encoding; Xenopus eFGF, Xenopus Smad2C (XSmad2C), human *Nanog* and human DPY30.

**Table 1:**
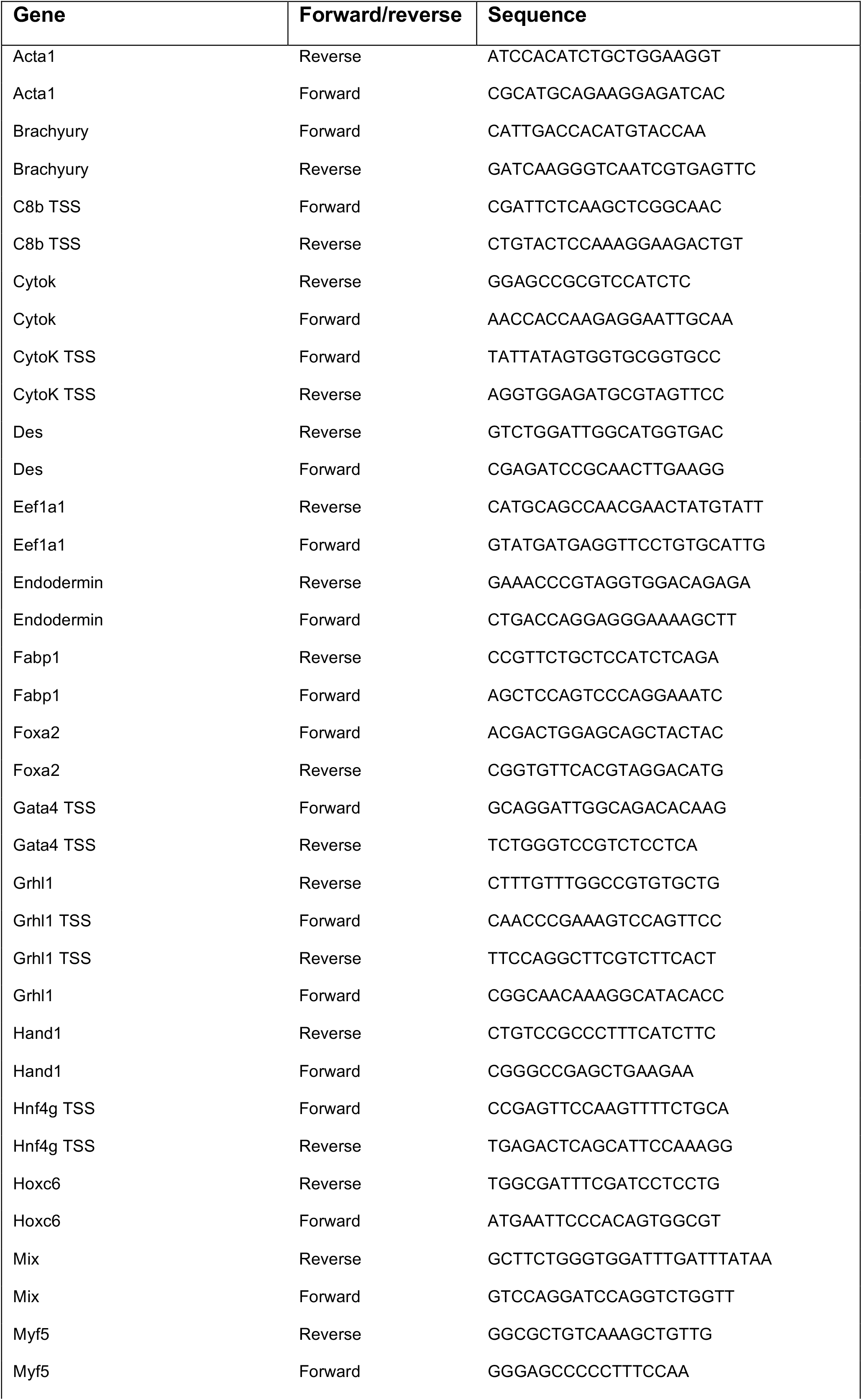
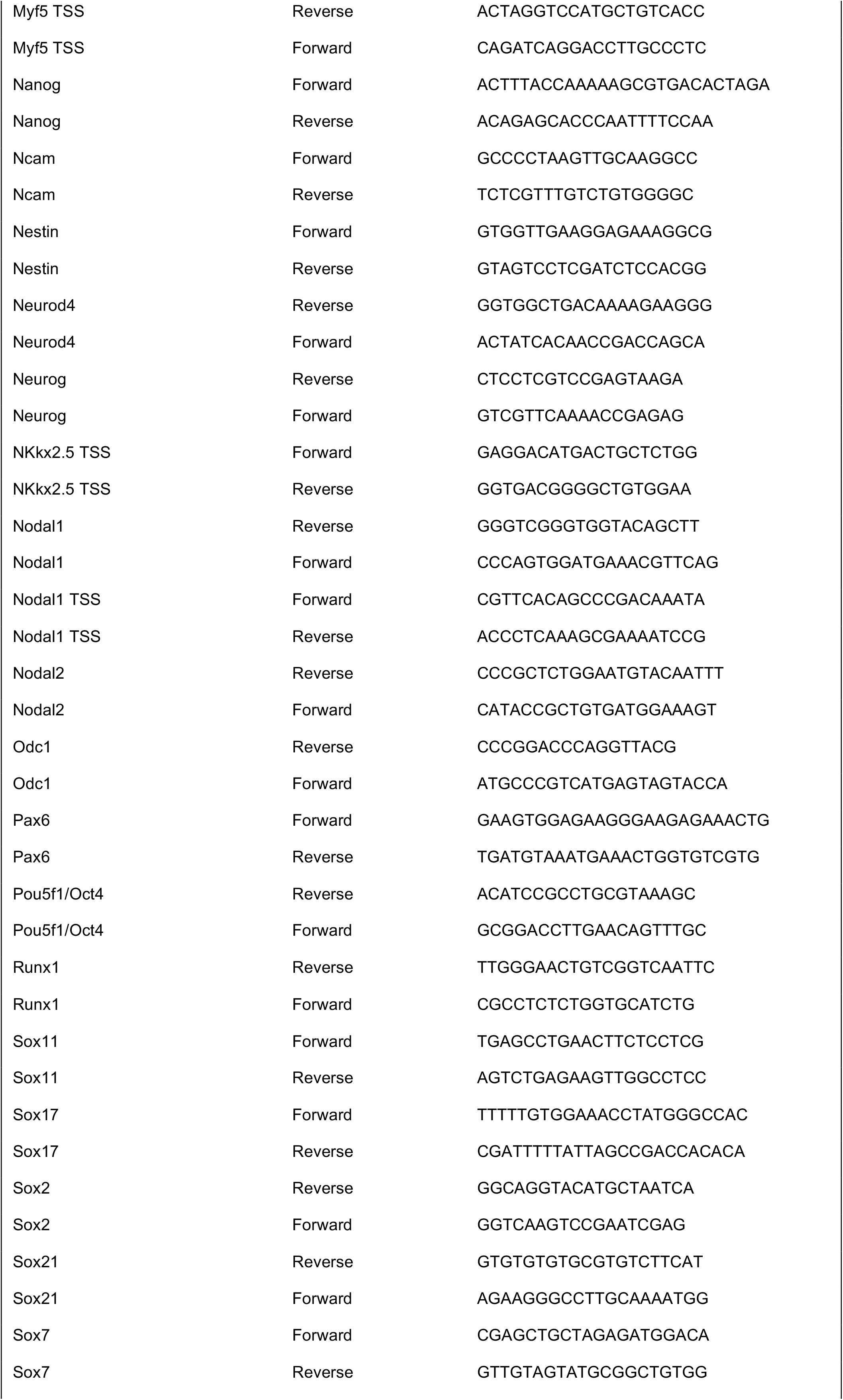
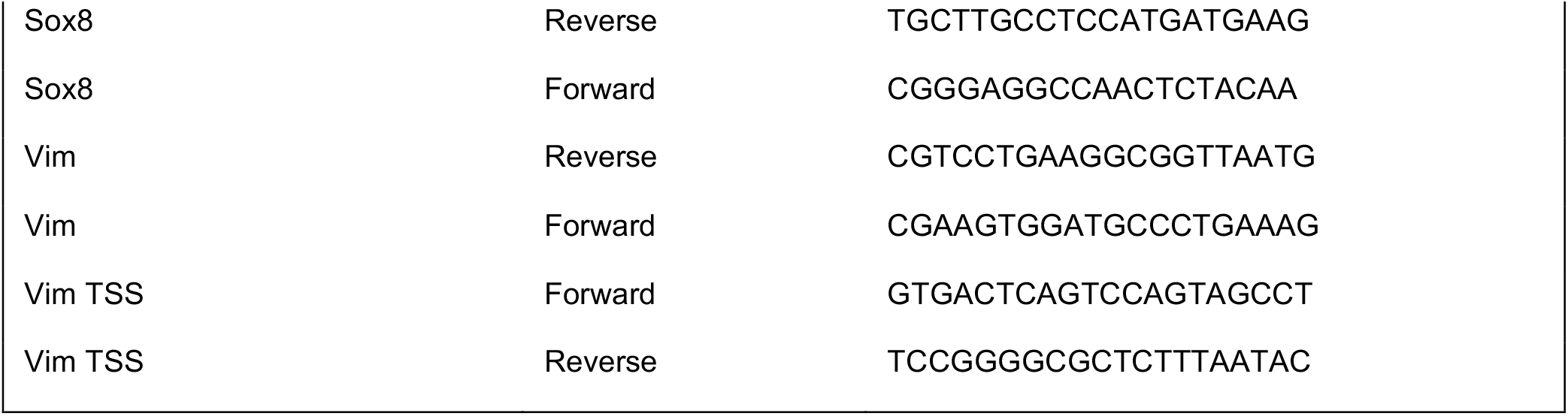
Primers used in this study.

### High-resolution electron microscopy (HREM)

Samples were fixed overnight at 4°C in 4% PFA in PBS before being dehydrated overnight in increasing concentrations of methanol (50%, 70%, 80%, 90% and 100%) before mounted in JB4 medium with acridine orange (SIGMA), and processed for HREM as described by Mohun and Weninger^27^. Section TIFF images were trimmed using the image processor function on photoshop before being imported into Osirix MD version 12.5.0 to produce embryo 3D reconstructions using the magma shader.

### Western blotting

For Western blot analysis, whole embryos were lysed in RIPA buffer and homogenised using a dounce homogeniser. Cell lysates were then centrifuged at 13,000 rpm at 4°C for 5 minutes. Supernatant was then collected, and protein concentration was assessed using Protein Assay reagent (BIORAD) according to manufacturer’s instructions. Approximately 25 µg of protein was used for each well for subsequent SDS-PAGE performed using standards methods. To test antibodies raised against axolotl *Nanog*, synthetic poly-A RNA encoding *Nanog*-HA were injected into mature Xenopus oocytes, from which lysates were prepared for Western blotting. In addition, an uninjected Xenopus oocyte lysate and a multiple antigen containing cell lysate were also used as HA-negative and positive controls, respectively. Axolotl *Nanog* antibodies were produced by DundeeCell, and raised against antigenic peptides specific to Ax*Nanog*. All other antibodies are commercially available and listed in table 2 as well as the used concentrations.

**Table 2:** Primers used in this study.

| Antibody | Species | Supplier | Catalogue number |
| --- | --- | --- | --- |
| Anti-H3K4me3 | Rabbit | Abcam | Ab8580 |
| Anti-H3K27ac | Rabbit | Abcam | Ab4729 |
| Anti-H3K36me3 | Rabbit | Abcam | Ab9050 |
| AntiH3K27me3 | Rabbit | Abcam | Ab6002 |
| Anti-H3 | Rabbit | Abcam | Ab1791 |
| Anti-HA | Rabbit | Abcam | Ab9110 |
| Anti-Nanog (axolotl) | Rabbit | Dundee cell | Custom made (available on request) |
| Anti-PolII | Rabbit | Abcam | Ab5131 |

### Immunofluorescence

Prior to immunofluorescence, embryos were incubated with an increasing gradient of sucrose/PBS solution (30%, 50% 80% sucrose/PBS) prior to mounting in OCT compound and frozen at -80° overnight. The blocks were transferred to -20° 2 hours before sectioning. Antigen retrieval was performed by boiling the slides in 0.01M citrate buffer (pH 6.0) for 10min. Sections were permeabilized with 1% Triton X-100 in PBS for 15min. Slides were then immersed in blocking solution (PBS supplemented with 5% BSA for 2h. After blocking, sections were incubated with primary antibody overnight at 4°C in a humidified chamber. Slides were then washed three times with 0.1% Tween-20/PBS before being incubated with a fluorescent secondary antibody (Extended Data table S2) for 45min at room temperature. Slides were treated with mounted with Fluoroshield with DAPI (Sigma) and sealed with nail varnish. Slides were kept at −20°C until observed.

### First-Strand cDNA synthesis was used to prepare samples for qPCR analysis

A reaction consisting of 1ul of Oligo(dT)20 (50uM), 2ug of total RNA, and 1ul of 10mM dNTP mix, was topped up to 13ul using sterile distilled water. The reaction was heated to 65° C for 5 minutes, and then incubated at 4°C for 5 minutes. The tube was briefly centrifuged before the addition of 4ul 5X First-Strand Buffer, 1ul 0.1M DTT, 1ul RNaseOUT recombinant RNase inhibitor (40 units/ul), and 1ul SuperScript III RT (200 units/ul). All reagents from Thermo Fisher Scientific. The reaction mix was then heated to 25°C for 5 minutes, 50°C for 60 minutes, and 70°C for 15 minutes, before being chilled to 4°C.

Each qPCR reaction contained 5ul of SYBR-Green Jumpstart Taq readyMix (SIGMA), 1ul (10uM [final]) of each of the forward and reverse primers (SIGMA), 2μl of nuclease-free water, and 1μl of cDNA template. Each reaction was prepared in triplicate on an ABI FAST Systems 0.2ml 96-well PCR plate (STARLAB), and then sealed with an Optical Adhesive cover (Life Technologies).

The following qPCR conditions were utilized for each run on the QuantStudio 6 Flex (Life Technologies) qPCR instrument: The plate was heated to 105°C to activate the Jumpstart Taq before being held at 50°C for 2 minutes. The initial denaturation step was at 94°C for 10 minutes, followed by 40 cycles of 94 °C for 15 seconds (denaturation) and 60 °C for 1 minute (annealing and extension). The raw data was then extracted to be analysed by comparative CT (Cycle threshold). An endogenous control gene was chosen for each qPCR run to normalise the data in order to compare relative fold change of target genes amongst cDNA samples. The double delta CT value (ΔΔCT) was calculated using the following formula: ΔCT target-ΔCT reference= ΔΔCT. As all calculations are in logarithm base 2, the expression fold change of each target gene was calculated using 2^-ΔΔCt.

### RNA-seq analysis

For staged Uninjected, Nanog KD, SB43 treated and DPY30 KD axolotl samples, RNA-seq, each sample was mapped to the transcriptome assembly described previously^25^ using RSEM with default parameters. This assembly was used as it also contains several developmental genes identified by our group in previous publications. Two of the Nanog KD stage 22 biological replicates were removed based on their lack of correlation with all other samples. The resulting count files were input to edgeR to calculate differentially expressed genes (FDR < 0.05, logFC > 1). Heatmaps were drawn using the log2 TPM values in R using heatmap.3 with default cluster settings.

The seventeen stages of wild-type axolotl early development was acquired by mapping the reads from the Jiang et al (2017) datasets^15^ to the same transcriptome assembly as before. Mean TPM values for each stage were calculated in R. These values were then used to determine whether each gene was ‘early’, ‘late’ or ‘global’. Genes that were only expressed with a TPM > 10 before stage 12 were classed as ‘early’, genes that were only expressed with a TPM > 10 on or after stage 12 were classed as ‘late’. The remaining genes were all considered global and were divided into ‘global high and ‘global low based on whether they had a TPM > 10 in every stage of the Jiang data or not. In order to construct the heatmap displaying every sample condition vs every sample condition, the heatmap row dendrogram was split at a height of 21, and each cluster was assigned the most common expression pattern of the genes within that cluster.

### Comparison of Human, Pig, Xenopus and tissues

In total, single-cell transcriptomes from 144 pig peri-gastrulation stage epiblast cells from Ramos-Ibeas and colleagues^7^, 152 human peri-gastrulation stage epiblast cells retrieved from Petropoulos *et al*^8^ were merged by calculating the geometric mean across all cells to get the simulated average count for each pluripotent tissue. This was then compared with whole RNA-seq data from st10.5 Xenopus animal caps retrieved from Angerilli *et al*^*6*^ and stage 10.5 axolotl animal caps (our study). Given the differences in sequencing methodology and sample preparation etc, we employed quantile normalisation to compare the relative abundance of expressed genes.

### Gene set enrichment using Xenopus cell type markers

The R package hypeR was used^28^ to perform a custom gene-set enrichment analysis of differentially expressed genes using a list of amphibian cell type markers identified by Briggs *et al*^*18*^. Analysis was carried out using an FDR threshold of 0.01. Cell types that were significantly enriched in the dataset were plotted using GOchord in R.

### ChIP QPCR

50 AC explants per experimental condition were processed using the truCHIP Chromatin Shearing Tissue Kit with Formaldehyde (Covaris). Caps were fixed with 1% methanol-free formaldehyde-PBS (Covaris) for 15 minutes before quenching. Fixed caps were processed according to the manufacturer’s protocols. 1ml of chromatin was sheared using the S220 ultra-sonicator (Covaris) in AFA fibre containing vesicles (Covaris). Chromatin shearing was performed under the following conditions: duty cycle 5%, intensity level 4, cycles/burst 200. The shearing program was run for 15 minutes resulting in chromatin fragments of 100–500 bp. 100µl of sheared chromatin was used per ChIP using the EZ MAGNA ChIP kit (Millipore) according to the manufacturer’s instructions. Purified DNA was then used for subsequent qPCR analysis. QPCR primers are listed in table 1.

## Data availability

All data generated or analysed during this study are included in this published article and its supplementary information files. All sequence data are deposited in the European Nucleotide Archive under the project number: PRJEB52201.

---

**Extended data Figure 1.**
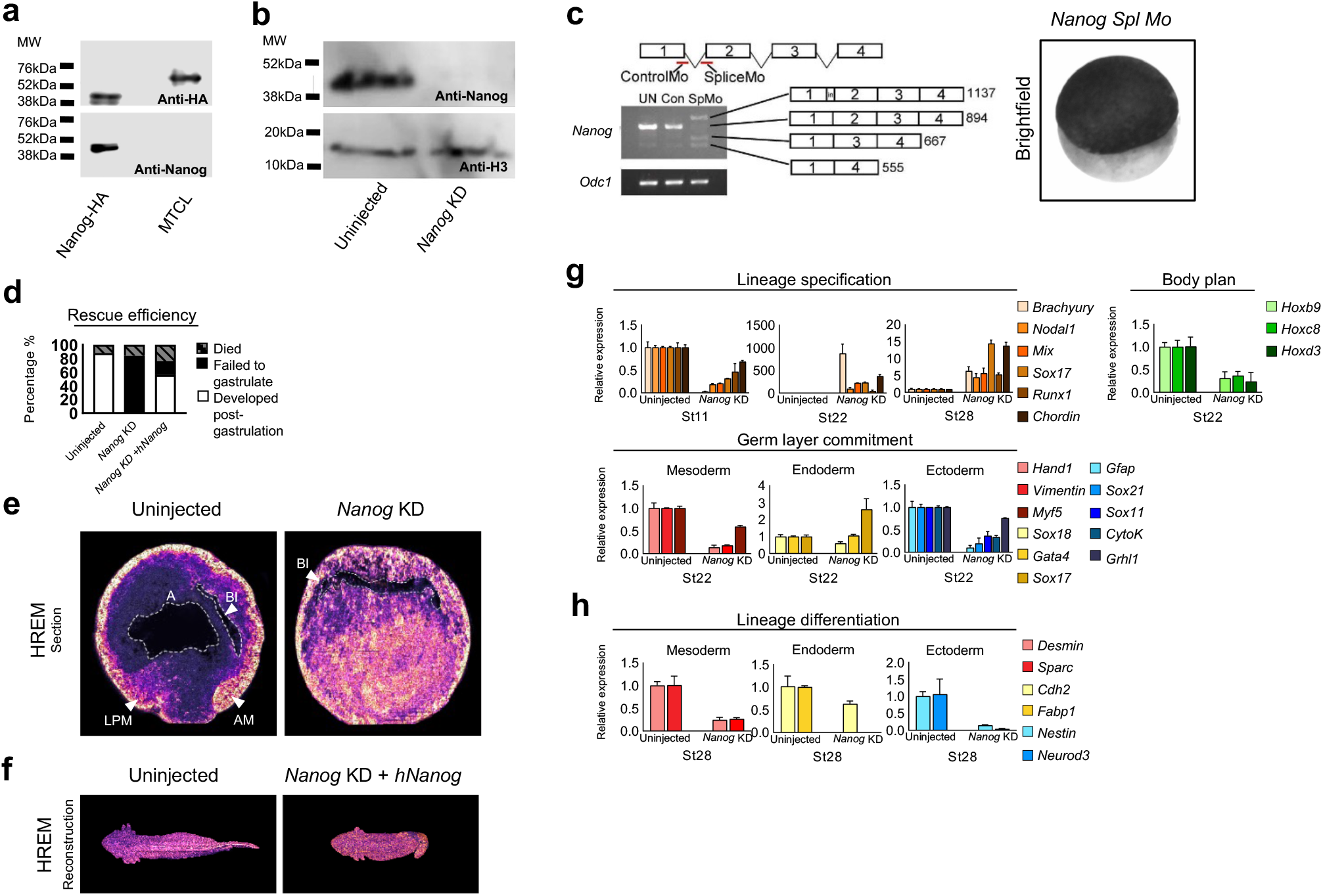
Nanog KD causes aberrant differentiation. **a**, Validation of antibodies raised against axolotl Nanog using western blotting. Lanes were loaded with lysates made from mature Xenopus oocytes following injection with Synthetic poly-A RNA encoding Nanog-HA or a multi-antigen tagged cell lysate. **b**, Western blot confirming complete KD of Nanog following MO injection. **c**, Validation of aberrant splicing in response to Nanog splice-morpholino and brightfield image of a stage 22 embryo following an injection of 80ng Nanog splice MO at the 1 cell stage (n=15). **d**, Nanog KD and rescue efficiencies (n=25). **e**, HREM imaging of mid-gastrula embryos with and without Nanog KD at stage 10.5. Visible structures highlighted: Involuting axial mesoderm (AM), ingressing ventral mesoderm (VM), archenteron (A), blastocoel (B). Scale bar, 1mm (n=2). **f**, HREM imaging of st42 uninjected and Nanog KD + hNanog rescue (dorsal view) (n=2). **g**, QPCR validation of key transcriptome findings (n=10). **h**, Differentiation markers of uninjected and Nanog depleted embryos at stage 28 (n=10× 3).

**Extended data Figure 2.**
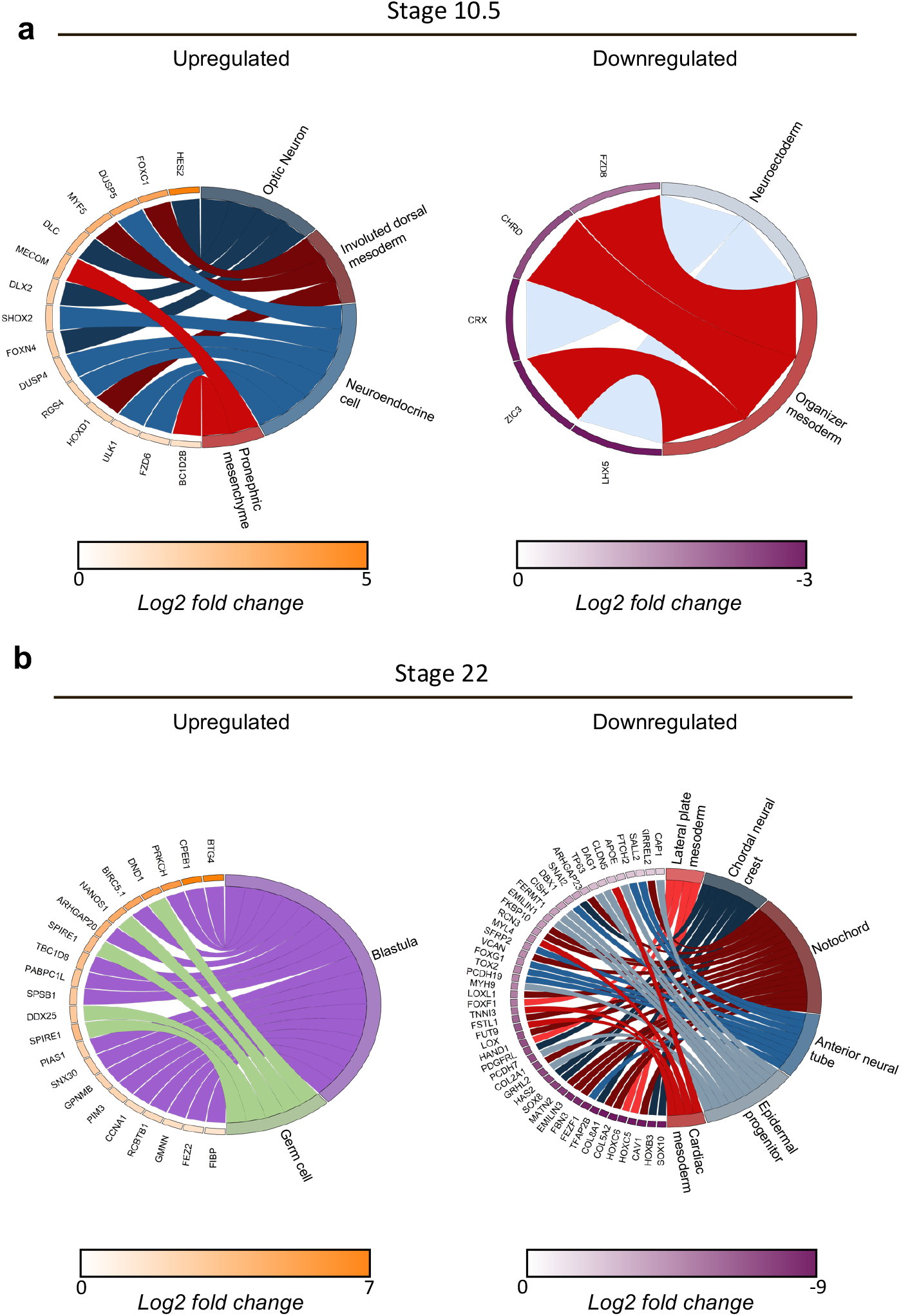
Enrichment of amphibian cell-type markers in Nanog KD DEGs. Chord diagrams showing the results of gene set enrichment analysis (GSEA) of amphibian cell-type specific markers in differentially expressed genes in *Nanog* KD. **a**, Nanog morphant upregulated DEGs are enriched for markers of optic neuron, involuted dorsal mesoderm, pronephric mesenchyme and neuroendocrine cells at stage 10.5. Downregulated DEGs at stage 10.5 are enriched for markers of neuroectoderm and organiser mesoderm. **b**, Upregulated DEGs in Nanog KD embryos stage 22 are enriched for markers of the blastula stage and germ cells. Downregulated DEGs are enriched for markers of lateral plate, notochord and cardiac mesoderm as well as markers of chordal neural crest cells, anterior neural tube and epidermal progenitor cells.

**Extended Data Figure 3.**
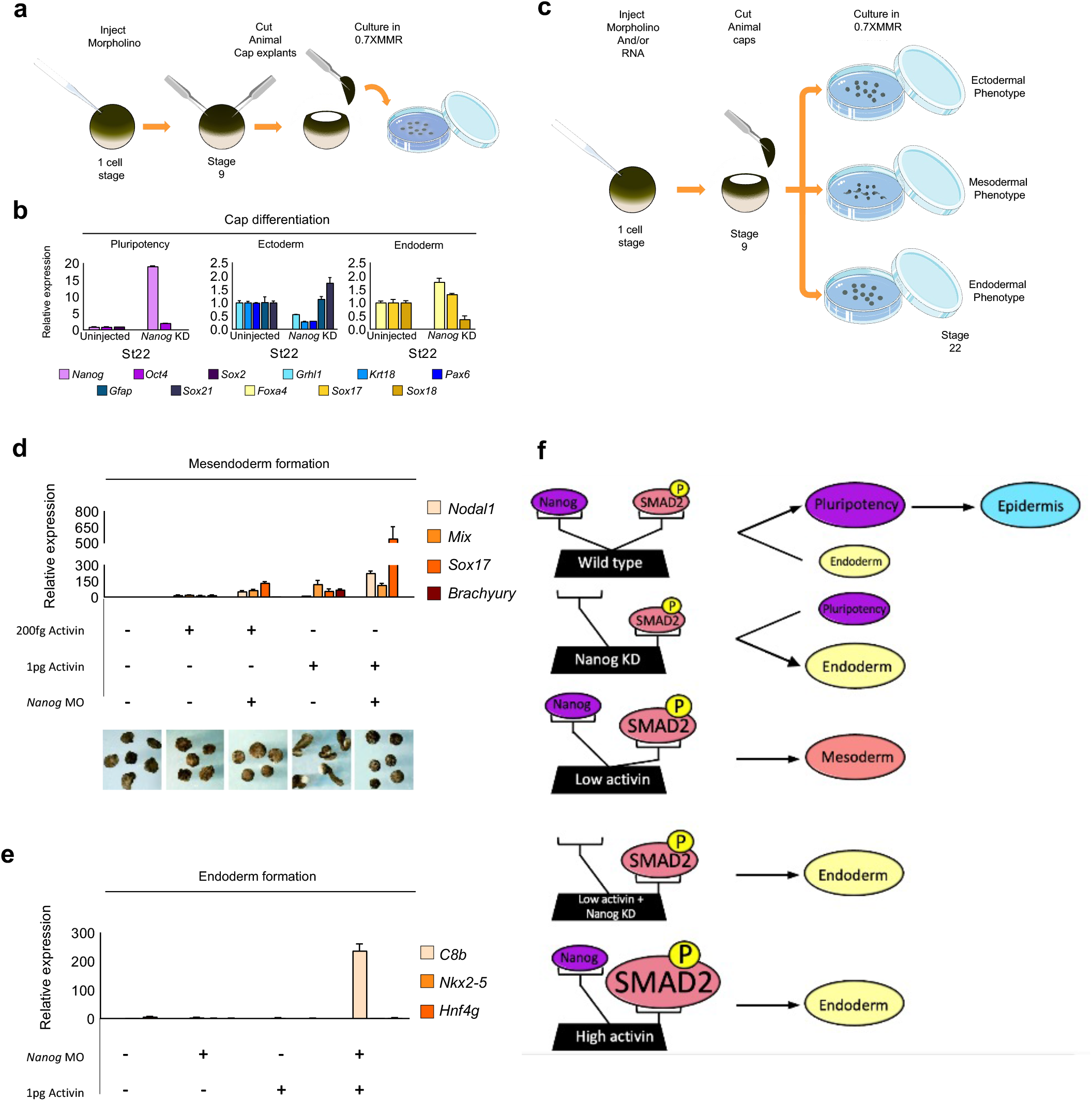
Nanog acts as a rheostat of SMAD2 activity. **a**, Schematic of morpholino injection regime and animal cap assay. **b**, Differentiation markers of uninjected and Nanog depleted AC explants at stage 22 (n=15). **c**, Schematic of mesoderm induction animal cap assay. **d**, Nanog KD increases activin sensitivity and prevents mesodermal but not endodermal differentiation in response to activin. QPCR of germ-layer markers of uninjected and Nanog depleted stage 22 caps following treatment with different activin concentrations (n=15). **e**, QPCR showing Nanog depleted caps express foregut/hindgut markers but not mature foregut/hindgut markers in response to activin (n=15). **f**, Schematic: Nanog acts as a rheostat of SMAD2 activity.

**Extended Data Figure 4.**
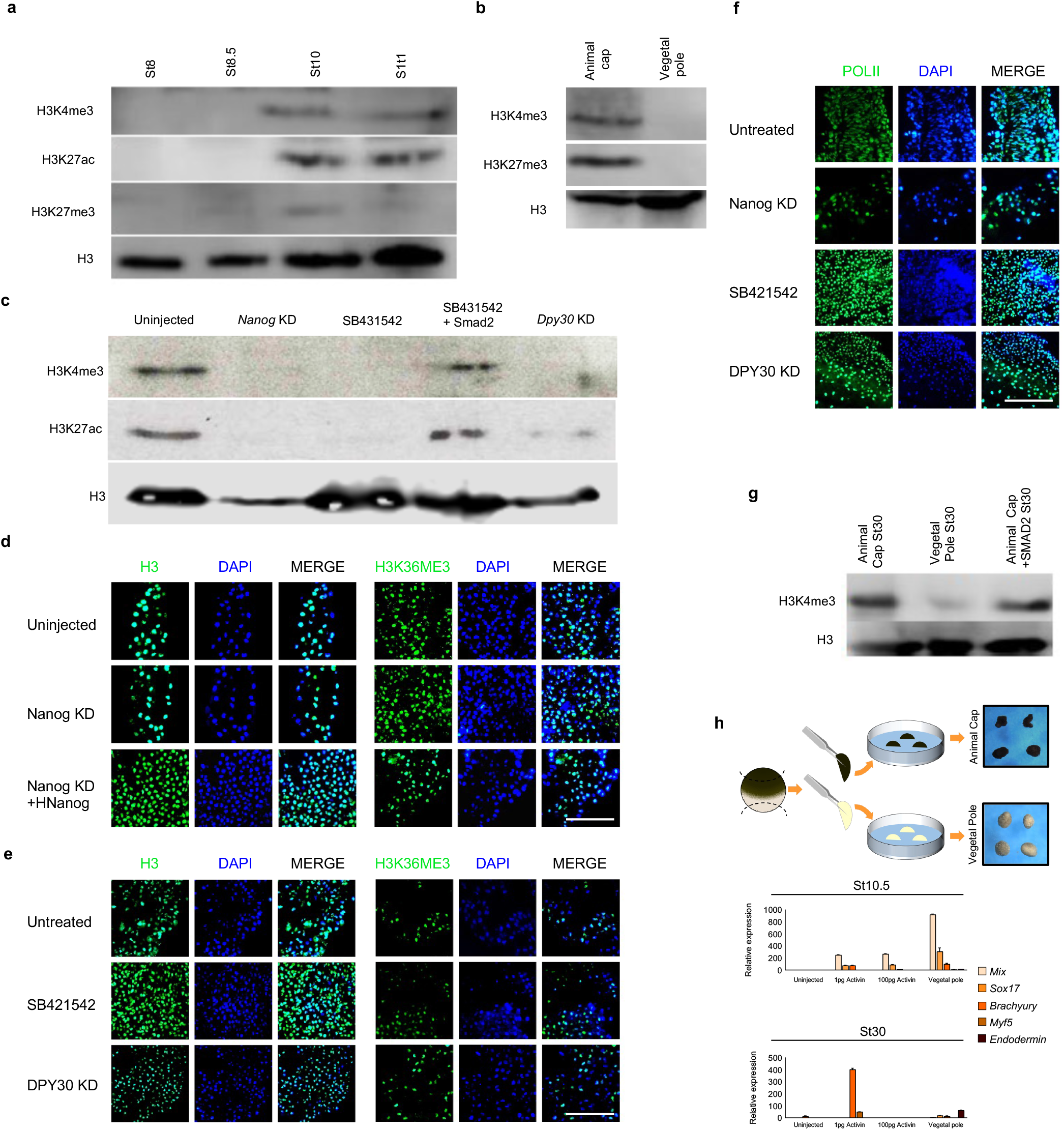
Nanog KD results in the loss of markers of activating promoter marks. **a**, Western blots showing H3K4me3, H3K27ac and H3K27me3 appear after ZGA at stage 8.5. **b**, Western blots showing vegetal explants have lower levels of H3K4me3. **c**, Western blotting confirming the depletion of H3K4me3 and H3K27ac in response to Nanog KD, SB431542 and DPY30 KD. **d**, Uninjected, Nanog depleted and hNanog rescued AC explants cultured to equivalent stage 14 and stained for H3, H3K36me3 and DAPI (n=3). Scale bar, 60µm. **e**, Untreated, SB431542 treated and DPY30 depleted animal cap explants cultured to equivalent stage 14 and stained for H3, H3K36me3 and DAPI (n=3). Scale bar, 60µm. **f**, Untreated, Nanog depleted, SB431542 treated and DPY30 depleted animal cap explants cultured to equivalent stage 14 and stained for phospho-POLII and DAPI (n=3). Scale bar, 60µm. **g**, Western blots showing vegetal explants show increased H3K4me3 in response to activin **h**, Vegetal explants form endoderm in a cell autonomous manner. QPCR of germ-layer markers of animal and vegetal explants compared with animal cap explants treated with different activin concentrations (n=15).

**Extended Data Figure 5.**
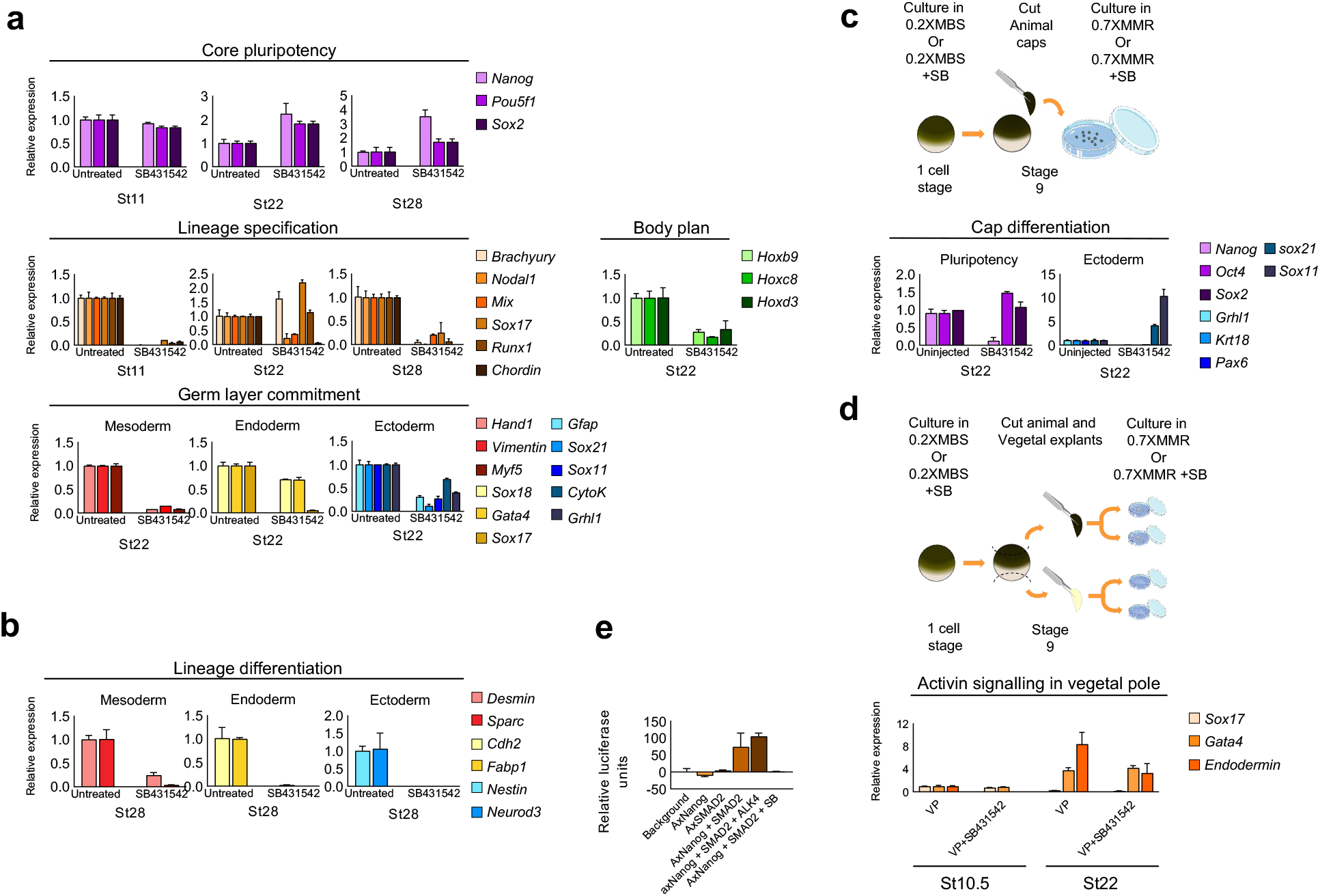
Validation of SB431542 transcriptome. **a**, QPCR validation of key transcriptome findings. **b**, QPCR analysis of late differentiation markers (n=15× 3). **c**, Differentiation marker expression of uninjected and SB431542 treated AC explants at stage 22 (n=15). **d**, Vegetal explants can form definitive endoderm even in the presence of SB431542 inhibitor. Vegetal explants from untreated or SB431542 treated embryos were cultured with or without SB431542 and assayed for endodermal markers (n=15). **e**, Luciferase complementation assay (n=3). Axolotl Nanog can physically interact with axolotl SMAD2, interactions are increased in the presence of constitutively active ALK4, binding is disrupted with SB431542 treatment.

**Extended Data Figure 6.**
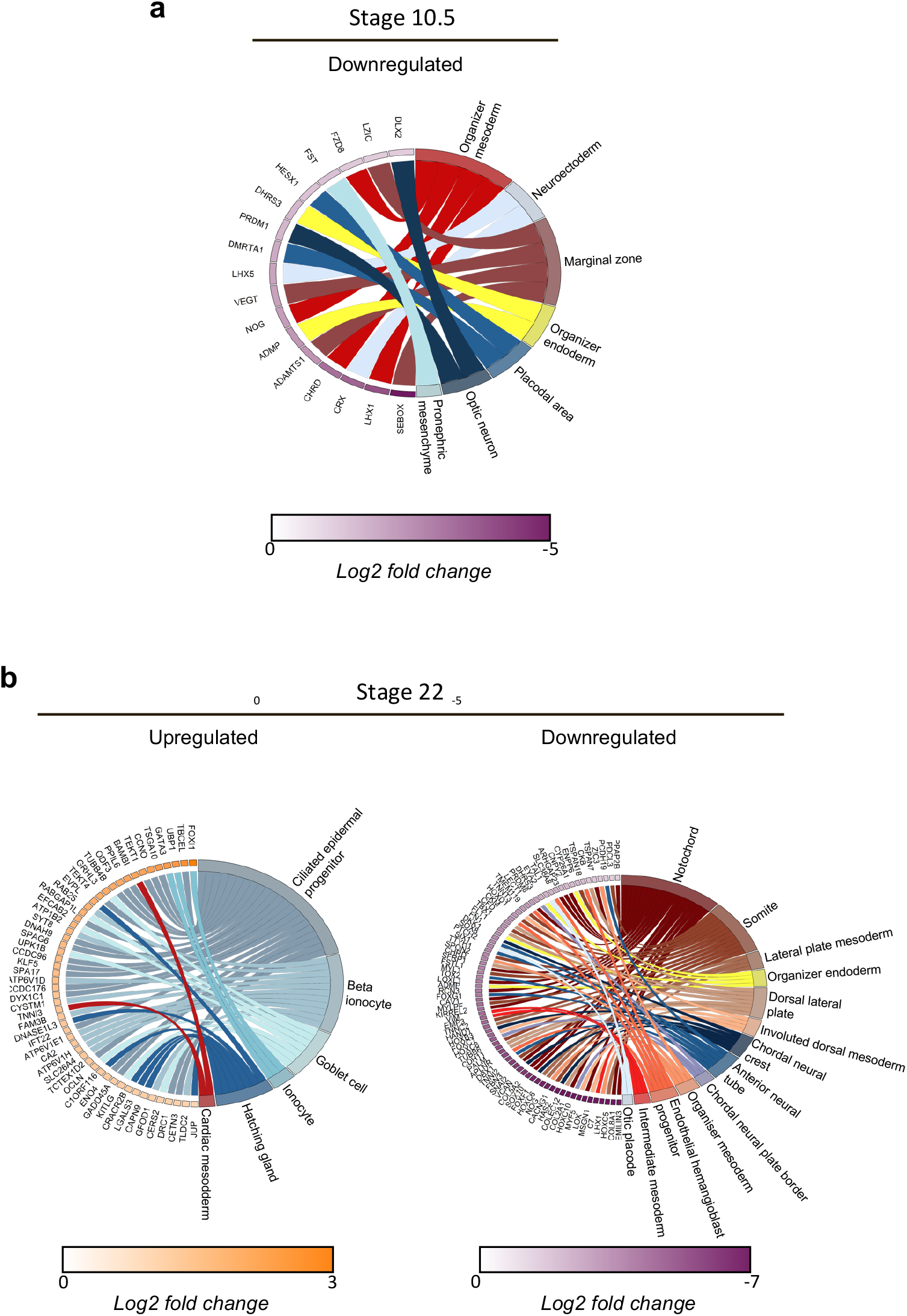
Enrichment of amphibian cell-type markers in SB43 DEGs. **a**, Chord diagrams showing the results of gene set enrichment analysis (GSEA) of amphibian cell-type specific markers in differentially expressed genes following SB43 treatment. **a**, SB treated embryo downregulated DEGs are enriched for markers of organiser and marginal zone mesoderm as well as organiser endoderm, neuroectoderm, placodal area, optic neuron and pronephric mesenchyme at stage 10.5. **b**, Upregulated DEGs in Nanog KD embryos stage 22 are enriched for markers of the ciliated epidermal progenitors, beta ionocytes, goblet cells, ionocytes, hatching gland cells and cardiac mesoderm. Downregulated DEGs are enriched for markers of notochord, somitic, lateral plate, dorsal, intermediate and organiser mesoderm as well as markers of organiser endoderm, chordal neural crest cells, anterior neural tube, otic placode and endothelial hemangioblast cells.

**Extended Data Figure 7.**
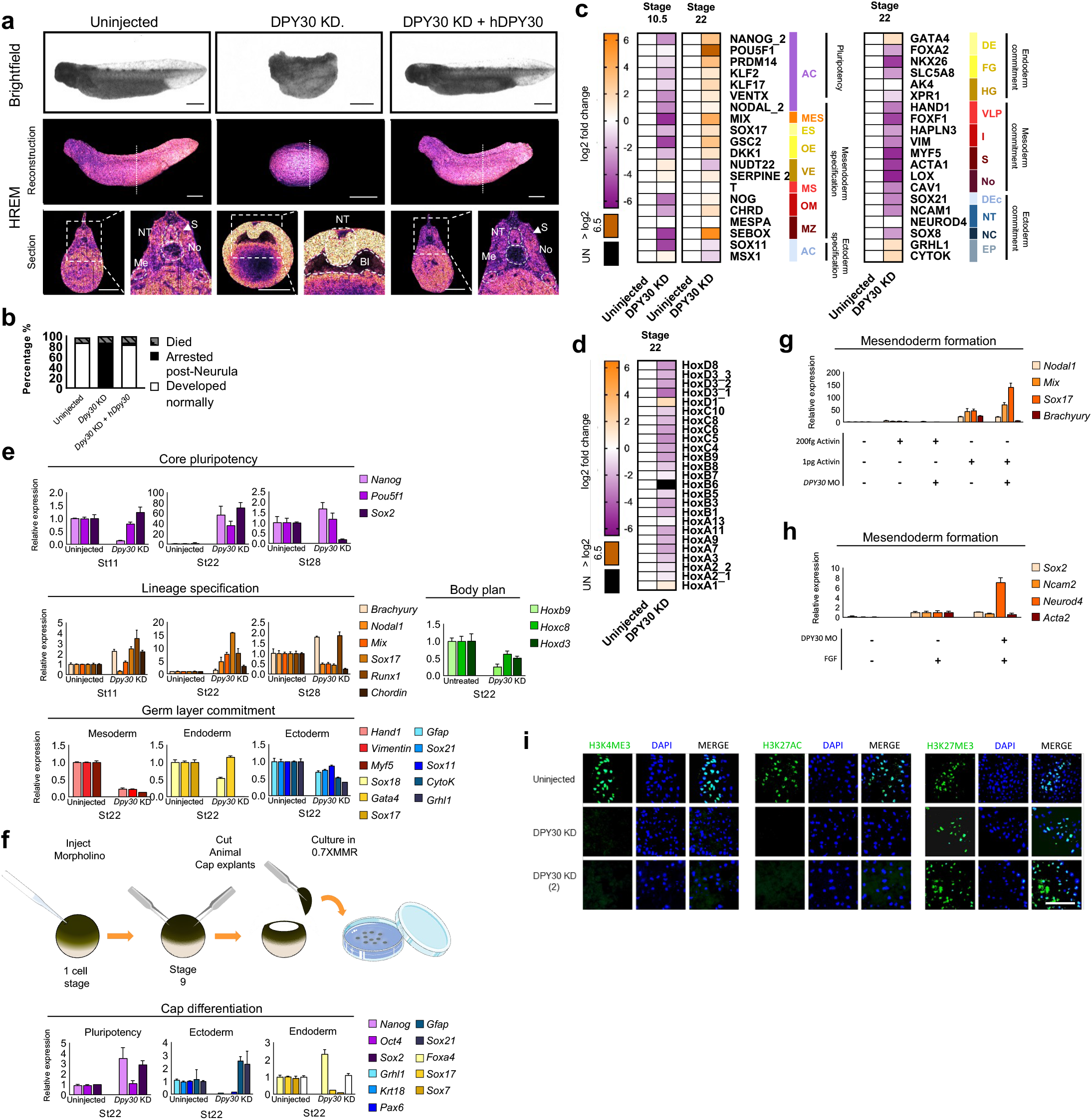
DPY30 KD resembles Nanog KD. **a**, DPY30 KD arrests development post-gastruation. Brightfield and HREM images of uninjected and DPY30 depleted embryos. Dotted line marks plane of section reconstruction. Dashed line delimits visible structures: Somites (S), Neural tube (NT) Notochord (No), Mesonephric ducts (Me), Blastocoel (B) (n=2). Scale bar, 1mm. **b**, DPY30 KD and rescue efficiencies (n= 30). **c**, Differential gene expression of key cell-type marker genes at stages 10.5 and 22 in uninjected (n=7 and 6, respectively) and DPY30 KD (n=6 and 6, respectively) embryos. Black indicates no detectable expression. Cell types: animal cap (AC), mesendoderm specification (general) (MES), endoderm specification (general) (ES), organiser endoderm (OE), vegetal endoderm (VE), mesoderm specification (general) (MS), organiser mesoderm (OM), marginal zone (MZ), definitive endoderm (general) (DE), foregut (FG), hindgut (HG), ventral-lateral plate (VLP), intermediate mesoderm (I), somite (S), notochord (No), definitive ectoderm (general) (Dec), neural tube (NT), neural crest (NC), epidermal progenitors (EP). **d**, Differential gene expression of Hox gene family members in DPY30 KD embryos at stage 22. **e**, QPCR of genes representative of key lineages. **f**, Differentiation markers of uninjected and Nanog depleted AC explants at stage 22 (n=15). **g**, DPY30 depletion prevents mesodermal but not endodermal differentiation in response to activin. QPCR of germ-layer markers of uninjected and DPY30 depleted stage 20 caps following treatment with different activin concentrations (n=15). **h**, DPY30 depletion reduces mesodermal gene expression and increased neuronal gene expression in response to FGF. QPCR of germ-layer markers of uninjected and DPY30 depleted stage 20 caps following treatment with FGF (n=15). **i**, Uninjected, DPY30 depleted embryos and hDPY30 rescued animal cap explants cultured to equivalent stage 14 and stained for H3K4me3, H3K27ac, H3K27me3 and DAPI (n=3). Scale bar: 60µm. 12

**Extended Data Figure 8.**
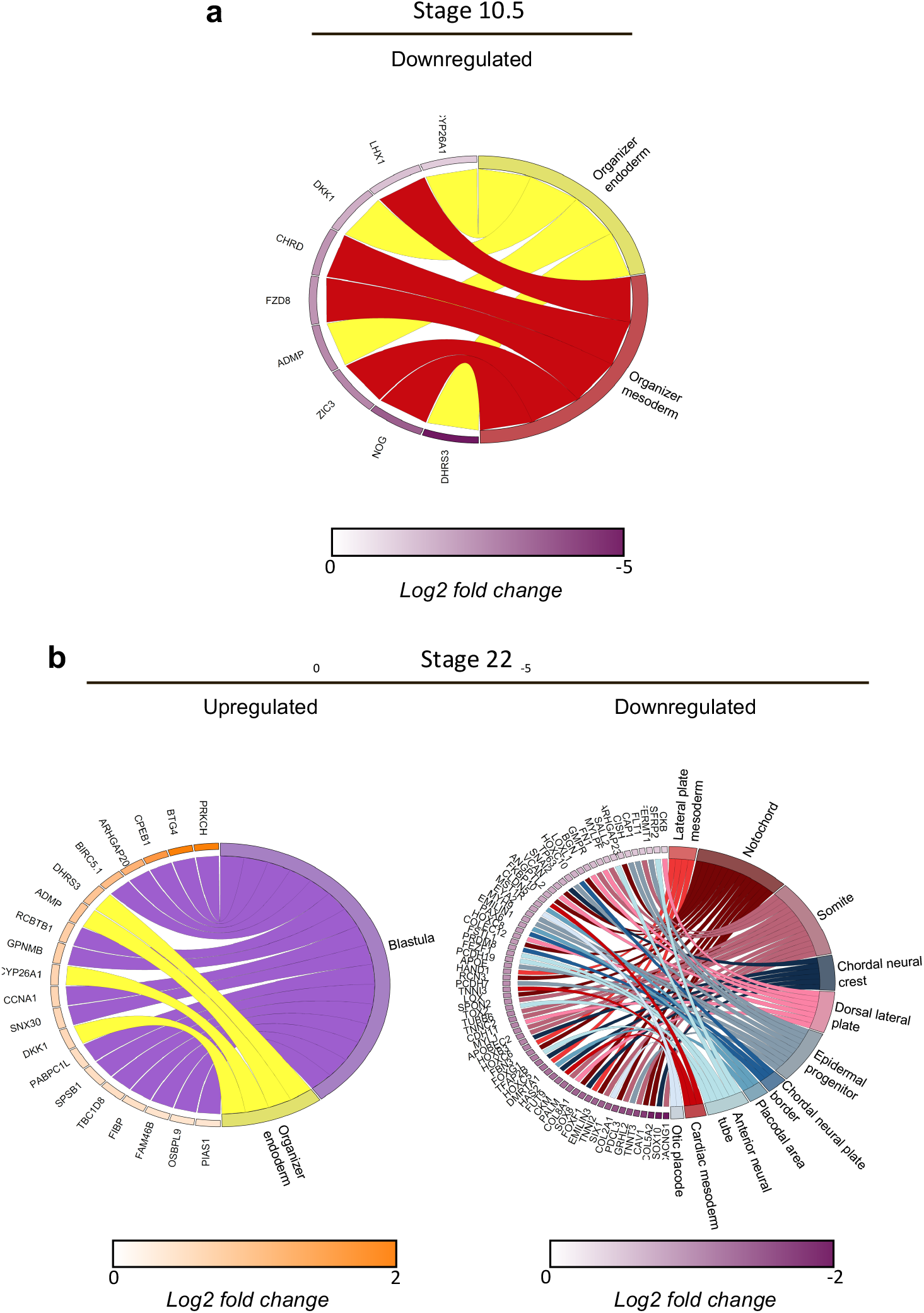
Enrichment of amphibian cell-type markers in DPY30 KD DEGs. Chord diagrams showing the results of gene set enrichment analysis (GSEA) of amphibian cell-type specific markers in differentially expressed genes following DPY30 KD. **a**, DPY30 morphant downregulated DEGs are enriched for markers of organiser mesoderm and endoderm at stage 10.5. **b**, Upregulated DEGs in DPY30 KD embryos stage 22 are enriched for markers of blastula cells and organiser endoderm. Downregulated DEGs are enriched for markers of lateral plate, notochord, somitic, dorsal leteral plate and cardiac mesoderm as well as markers of chordal neural crest cells, anterior neural tube, otic placode, chordal neural plate border, anterior neural tube and epidermal progenitor cells.

**Extended Data Figure 9.**
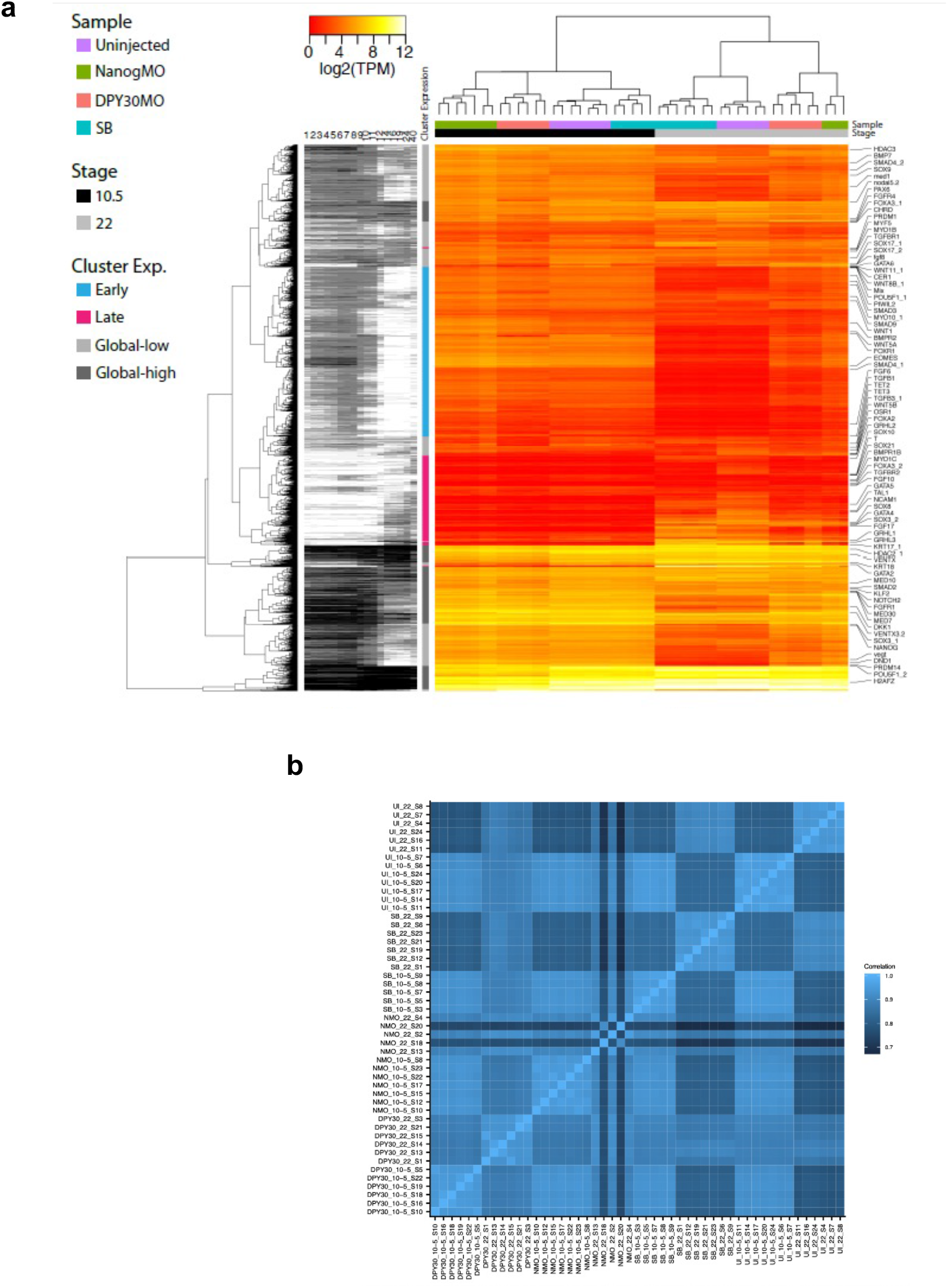
Comparison of Nanog KD, DPY30 KD and SB431542 treatment transcriptomes. **a**, Heatmap showing hierarchal clustering of ‘early’ or ‘late’ genes in uninjected, SB431542 treated, Nanog and DPY30 depleted embryo transcriptomes at stages 10.5 and 22. The ‘early’ genes were defined by genes which had an expression greater than 10TPM and had a significantly lower expression at stage 10.5 than at stage 22. Correspondingly, ‘late’ genes were defined by genes which had a TPM greater than 10 at stage 22 and had significantly lower expression at stage 10.5. Generally early activated genes are more highly expressed even in late stage embryos following SB treatment or Nanog/DPY30 KD. **b**, Heatmap showing the Pearson correlation of the whole transcriptome of each sample.

**Extended Data Figure 10.**
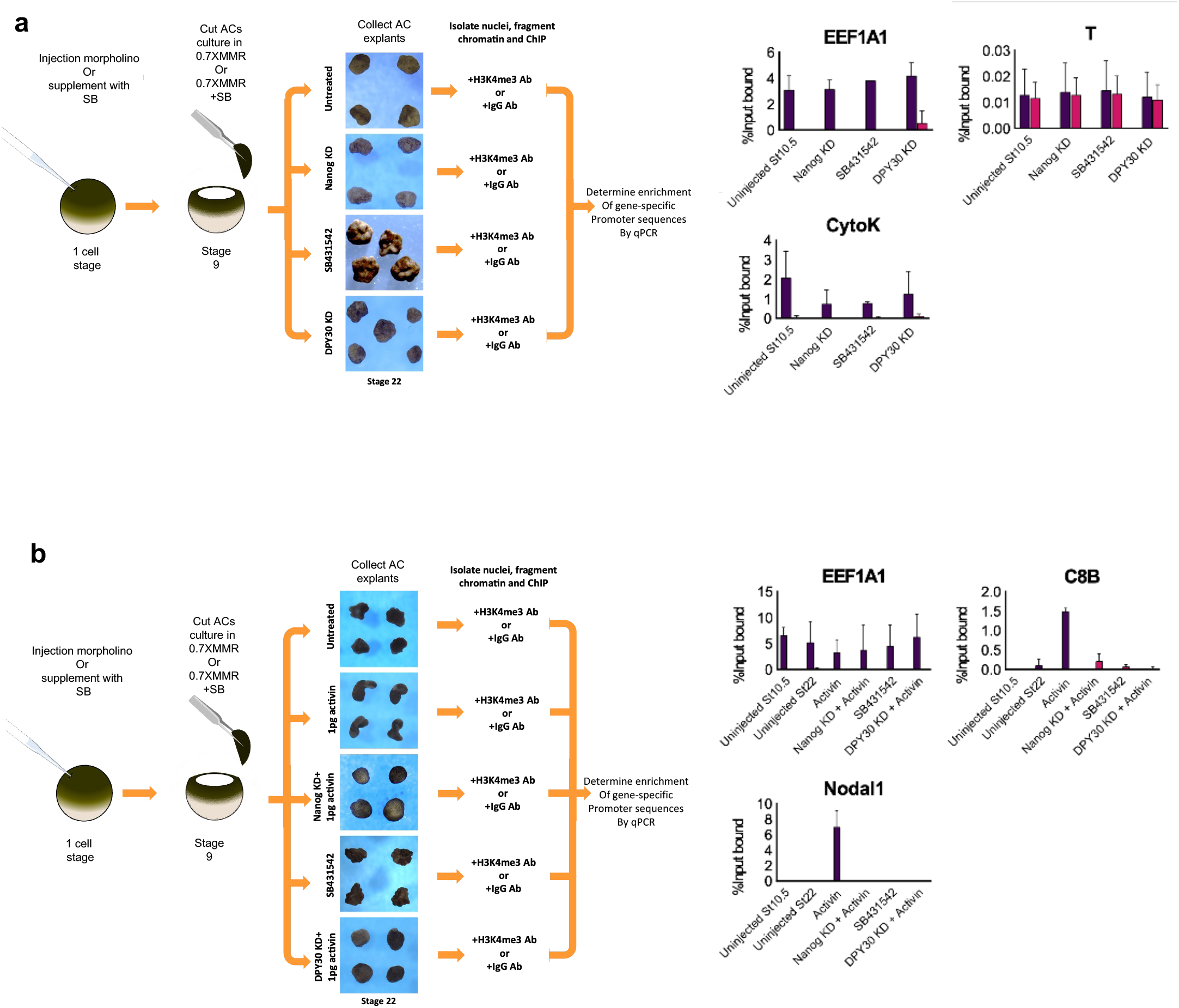
Nanog gene target loci lack H3K4me3. **a**, H3K4me3 ChIP of stage 10.5 uninjected, Nanog KD, DPY30 KD and SB431542 treated caps followed by qPCR using probes directed at gene promoter regions (n=50 per experimental condition). **b**, H3K4me3 ChIP-qPCR of equivalent stage 10.5 and 22 uninjected caps, stage 22 Activin treated caps with and without Nanog and DPY30 depletion as well as SB431542 treated caps (n=50 per experimental condition).

